# *ASXL1* truncating mutations drive leukemic resistance to T cell attack

**DOI:** 10.1101/2025.05.29.656798

**Authors:** Dustin McCurry, Zhongqi Ge, Jaehyun Lee, Rishi Pasumarthi, Xiaohong Leng, Thomas Koehnke, Oren Pasvolsky, Pranaya Raparla, Vinhkhang Nguyen, Katie Maurer, Shuqiang Li, Kenneth J. Livak, Emerie Danson, Bijal Thakkar, Elham Azizi, Robert J. Soiffer, Sachet A. Shukla, Ravi Majeti, Jerome Ritz, Catherine J Wu, Jeffrey J. Molldrem, Pavan Bachireddy

## Abstract

We previously found that specific exhausted T cell subsets defined response, but not resistance, to donor lymphocyte infusions (DLI), a curative immunotherapy for leukemic relapse following allogeneic stem cell transplant (SCT). To identify leukemia molecular pathways that drive resistance, we analyzed whole exome and targeted mutation panel sequencing in two independent cohorts of DLI-treated patients, nominating oncogenic, truncating mutations in *ASXL1* (*ASXL1*^MUT^) as the genetic basis for DLI resistance. Deep interrogation of 138,152 bone marrow single myeloid cell transcriptomes (scRNA-seq) from this cohort linked DLI resistance to a transcriptional state notable for leukemic stem cell identity and HLA-I suppression. *In silico* analysis of publicly available scRNA- and ATAC-seq data in acute myeloid and chronic myelomonocytic leukemias, respectively, confirmed an association between *ASXL1*^MUT^ and HLA-I suppression across myeloid malignancies. CRISPR correction of the endogenous *ASXL1*^MUT^ in the K562 leukemic cell line increased HLA-I, but not HLA-II, surface protein expression through increased deposition of the activating H3K4Me3 mark with only modest effect on the repressive H3K27Me3 mark, suggesting a Polycomb-independent mechanism of action. Indeed, inhibitors of EZH2, a critical component of the PRC2 complex, significantly upregulated HLA-I surface protein expression independently of *ASXL1*^MUT^, suggesting that EZH2 inhibition could bypass *ASXL1*^MUT^-mediated HLA-1 suppression. Importantly, *ASXL1*^CORRECTION^ significantly increased CD8+ T cell recognition, activation and killing, and *ASXL1*^MUT^-mediated T cell suppression could be overcome by EZH2 inhibition. Thus, by integrating molecular analyses with immuno-functional studies, we define a novel oncogene-driven pathway of immune evasion and propose a therapeutic strategy to re-engage T cell killing in *ASXL1*^MUT^ tumors.

## Main Text

The therapeutic graft-versus-leukemia (GvL) effect is critical to the clinical activity of allogeneic stem cell transplantation (allo-SCT), a potentially curative form of immunotherapy for patients with leukemia.^1–3^ GvL resistance (GvL-r) is a critical driver of morbidity and mortality following allo-SCT as outcomes following leukemic relapse are dismal.^3^ One approach that seeks to address GvLr is donor lymphocyte infusion (DLI),^3–6^ an adoptive immunotherapy with a spectrum of clinical activity. To probe the drivers of GvL-r in response to DLI, we previously characterized the T cell states in responders (R) or non-responders (NR).^7^ Strikingly, we identified that specific exhausted T cell subsets defined DLI responders (DLI-R); though we were unable to identify a pattern of T cell dysfunction in non-responders (DLI-NR)^7^ suggesting that some forms of GvL-r may be mediated by T cell extrinsic features. Thus, we hypothesized that leukemic molecular features may define resistance to DLI, informing our understanding of GvL-r.

To test this hypothesis, we assembled a cohort of 14 patients treated with CD8-depleted DLI for relapsed CML following allo-SCT. Eight patients were long-term DLI-R, defined as having achieved molecular remission (i.e., RT-PCR negative for the BCR-ABL transcript) after DLI, and six were DLI-NR, who did not achieve molecular remission following DLI. As published previously, none of the patients developed acute graft-versus-host disease (GvHD) after DLI and the timing between allo-SCT and DLI therapy was comparable between DLI-R and DLI-NR.^7^ As a reference, we also analyzed post-transplant BM biopsies from two patients with CML who never relapsed after allo-SCT, and as a confirmation cohort, we assembled an independent set of six patients (1 R and 5 NR) treated with CD8-depleted DLI for relapsed CML at a separate institution. In this independent cohort, one patient developed GvHD following DLI. Serial BM or PB samples were collected at a median of 3 time points per patient. Whole exome sequencing was performed on 49 samples from 11 patients within the discovery cohort. A targeted mutation panel^8^ was performed on seven samples from the independent cohort of six patients. For 46 samples, we obtained single-cell RNA sequencing (scRNA-seq) data on viable mononuclear cells. Additional cohorts consisted of single-cell chromatin accessibility sequencing (scATAC-seq) in five patients with chronic monomyelocytic leukemia (CMML)^9^ and nine patients with acute myeloid leukemia (AML)^10^ (Figure 1a).

**Figure 1.**
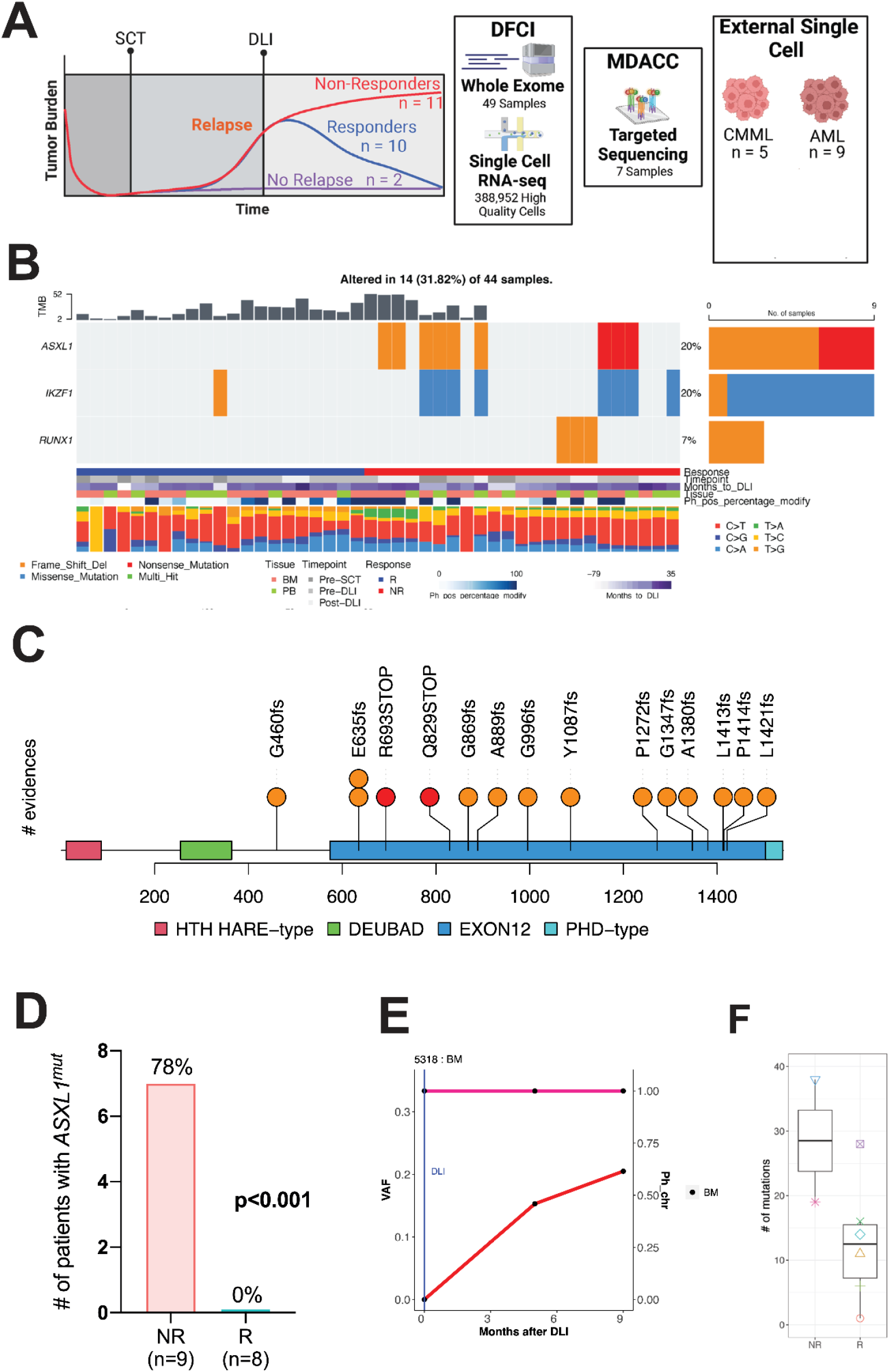
ASXL1 mutations are the genetic basis for DLI resistance. A) Cartoon schematic of molecular analyses and clinical cohorts. B) Co-mutation plot illustrating the recurrent cancer drivers from whole exome sequencing of pre-/post-SCT and pre-/post-DLI CML relapses. C) Lolliplot plot indicating predominance of *ASXL1* mutations in exon 12. D) Association of *ASXL1* mutations with DLI resistance in expanded cohort. E) Example of subclonal selection for *ASXL1* mutations during DLI resistance. F) Total mutation burden in pre-DLI samples from responders/nonresponders.

### Identification of *ASXL1* mutations in DLI resistance

Within the discovery cohort we obtained genomic DNA from recipient-derived mesenchymal stromal cells (R-MSC) and donor peripheral blood cells (D-PBC) for germline sources in eleven patients (7 R and 4 NR). To identify somatic alterations, we used MuTect2 with R-MSC and D-PBC as controls. Given the prognostic role of additional oncogenic driver mutations in CML, both in the pre- and post-TKI era,^11^ we sought to identify mutations in known leukemic driver genes. Strikingly, no recurrent mutations in known cancer drivers were identified at post-SCT relapse time points in patients that responded to DLI. We identified recurrent mutations in *ASXL1, IKZF1*, and *RUNX1* in 14 of 44 samples (31.82%), all from patients that did not respond to DLI (NR) (Figure 1b). Focusing on pre-DLI samples, known non-Philadelphia (Ph) chromosome cancer drivers were found in 4/4 DLI-NRs (*RUNX1* (1/4), *IKZF1* (2/4), and truncating exon 12 *ASXL1* mutations (*ASXL1*^m^) (3/4)). *ASXL1*, IKZF1, and RUNX1 mutations are linked to prognosis, advanced-phase, and myeloid and/or lymphoid skewing in CML.^11^ All IKZF1 mutations (2/4) were predominantly missense mutations, in contrast to nonsense or frameshift mutations more commonly reported in CML.^12^ Each patient with an IKZF1 mutation had a concurrent *ASXL1*^m^, and one additional DLI-NR was found to have an *ASXL1*^m^ only. One patient had a frame-shift mutation in RUNX1, a common mutation type in myeloid malignancies.^13^ All *ASXL1* mutations identified in 3/4 NR were frameshift or nonsense mutations within exon 12 prior to the plant homeodomain (PHD), consistent with the commonly mutated region in myeloid malignancies.^14–16^ These mutations are reported to produce a truncated protein with altered function. We sought to determine whether these findings were reproducible, so we performed targeted mutation sequencing^8^ on seven samples from an independent cohort of six CML patients (1 R and 5 NR) treated with CD8-depleted DLI at MD Anderson Cancer Center. We identified *ASXL1* mutations in 4/5 NR and 0/1 R. The most common mutation was E635fs (2 patients) (Figure 1c). In total, we identified *ASXL1* exon 12 nonsense or frame-shift mutations in 7/9 NR (78%) and 0/8 R (0%) (p=7e-4, two-sided Fisher’s test) (Figure 1d). In one patient, 5318, DLI heralded the emergence of mutant *ASXL1*, further rising over time, suggesting clonal evolution of the CML following DLI, despite the same Ph-chromosome positivity at all time points (Figure 1e).

Given the potential role for neoantigens to participate in the GvL response, we also calculated the tumor mutation burden (TMB),^17,18^ and sought to determine whether DLI response is associated with TMB or predicted neoantigens with strong binding affinity (defined as binding affinity ≤50 nM by netMHCpan). Our prior findings^17,18^ suggested that neoantigen specific responses are maintained in long-term responders to allo-SCT and depleted in patients who relapse late after allo-SCT. We did not identify either TMB or predicted high affinity neoantigen burden as predictive biomarkers for DLI response (Figure 1f). Two patients received greater than one allo-SCT prior to DLI, limiting the ability to calculate a total mutational burden for these patients.

### Identification of leukemic cells in scRNA-seq Data

The emergence of *ASXL1* mutations in NR led us to hypothesize that DLI resistance may be mediated by leukemic intrinsic features shaped by *ASXL1* mutations. To test this hypothesis, we interrogated our cohort of scRNA-seq samples. In total, this cohort was composed of 57 samples from 17 patients (8 R, 6 NR, and 2 patients who never relapsed). Following batch correction (Harmony) and quality filtering, we identified 388,952 cells passing QC (Figure 2a). Consistent with findings in other chronic myeloid malignancies,^19^ we identified intact hematopoietic lineages, including T-cells, B-cells, megakaryocyte-erythroid precursors (MEP), and monocytes (Figure 2a). To identify leukemic cells, we integrated our WES sequencing to identify expression of mutant transcripts (Figure 2b). Mutant cells were largely confined to the myeloid compartment. Mutant cells composed 0.19% of all cells, which was correlated with Ph+ burden (Figure 2b). Consistent with the known challenges of detecting *ASXL1* mutations in scRNA-seq data, we were unable to detect *ASXL1* mutant cells.^20,21^

**Figure 2.**
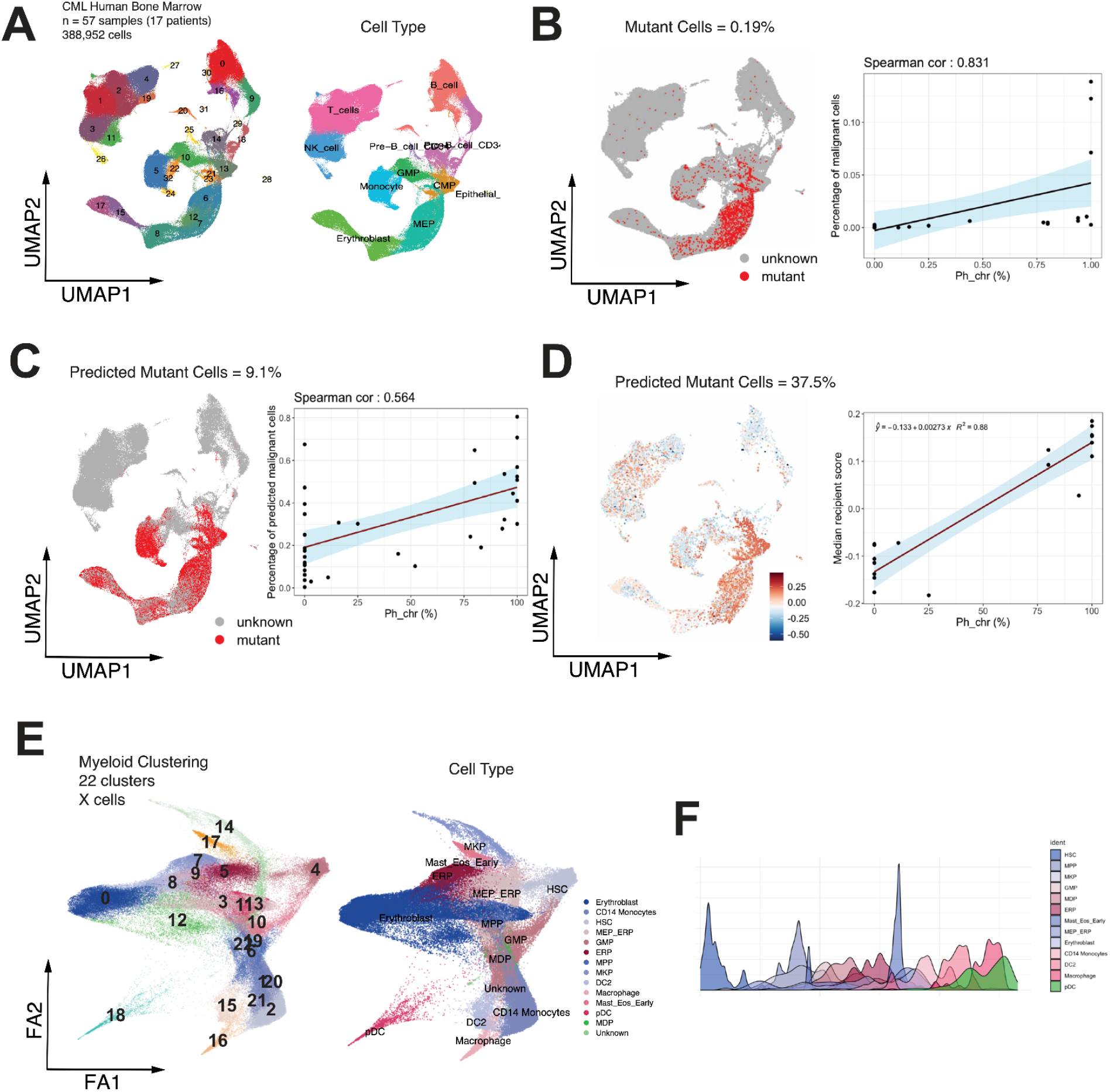
Single cell transcriptomic landscape of DLI therapy. A) UMAP clustering of all hematopoietic cells, with relevant cell types indicated. B) Cells with WES-derived somatic variants identified in transcriptomes colored in red. C) Results of training and prediction of mutant cells with ML classifier and correlation with %Ph positivity. D) Prediction of leukemic cell identity based instead on recipient-donor chimerism results in improved correlation with %Ph positivity. E) UMAP of myeloid cells after harmonization and batch correction. F) Cell states ordered across pseudotime.

Given the low frequency of mutant cells when integrating WES with scRNA-seq data, we attempted other approaches to identify leukemic cells to assess leukemic intrinsic features. A machine learning approach based upon transcriptional states of known mutant cells from exome data^22^ (Figure 2c) demonstrated over a ten-fold increase in mutant cell yield (9.1%), but was not well correlated with Ph+ burden (Figure 2c). In a subset of cases, alloSCT were performed with sex mismatched donors, enabling detection of Y-chromosome derived transcripts uniquely in the donor or recipient, enabling calculation of a donor-recipient score (D/R score). This measure demonstrated a significantly improved correlation with Ph+ burden, identifying 37.5% of mutant cells (Figure 2d).

### DLI-r links to suppressed HLA-I in LSC Compartment

Given the predominance of recipient cells within the myeloid compartment, we performed a reclustering of the myeloid compartment in Scanpy with a force directed layout^23^ to better assess the relationships between cell states (Figure 2e). To identify hematopoietic trajectories and transitions in cell state, Monocle 3 was used^24^ (Figure 2f). We identified a progenitor cell state and were able to recapitulate known hematopoietic differentiation trajectories, including erythroid, myeloid, and megakaryocytic lineages. Gene programs identified in differentiating hematopoietic progenitors identified in post stem-cell transplant bone marrow samples were used.^25^ Within this re-clustering, we continued to identify intact hematopoietic lineages consistent with our prior finding (Figure 2e,2f). To assess for cellular states that herald DLI resistance, we performed a differential enrichment analysis based on responders or nonresponders at timepoints immediately prior to DLI. This identified one cluster composed of monocytes enriched in R (Figure 3a), consistent with results in other cancer types treated with PD-1 immunotherapy.^26^ In nonresponders, we identified four progenitor clusters enriched in non-responders composed of granulocyte-monocyte progenitors (GMP), basophil and eosinophil progenitors (Eo/BPro), megakaryocyte progenitors (MkP), and a hematopoietic stem cell-like cluster (HSC).

**Figure 3.**
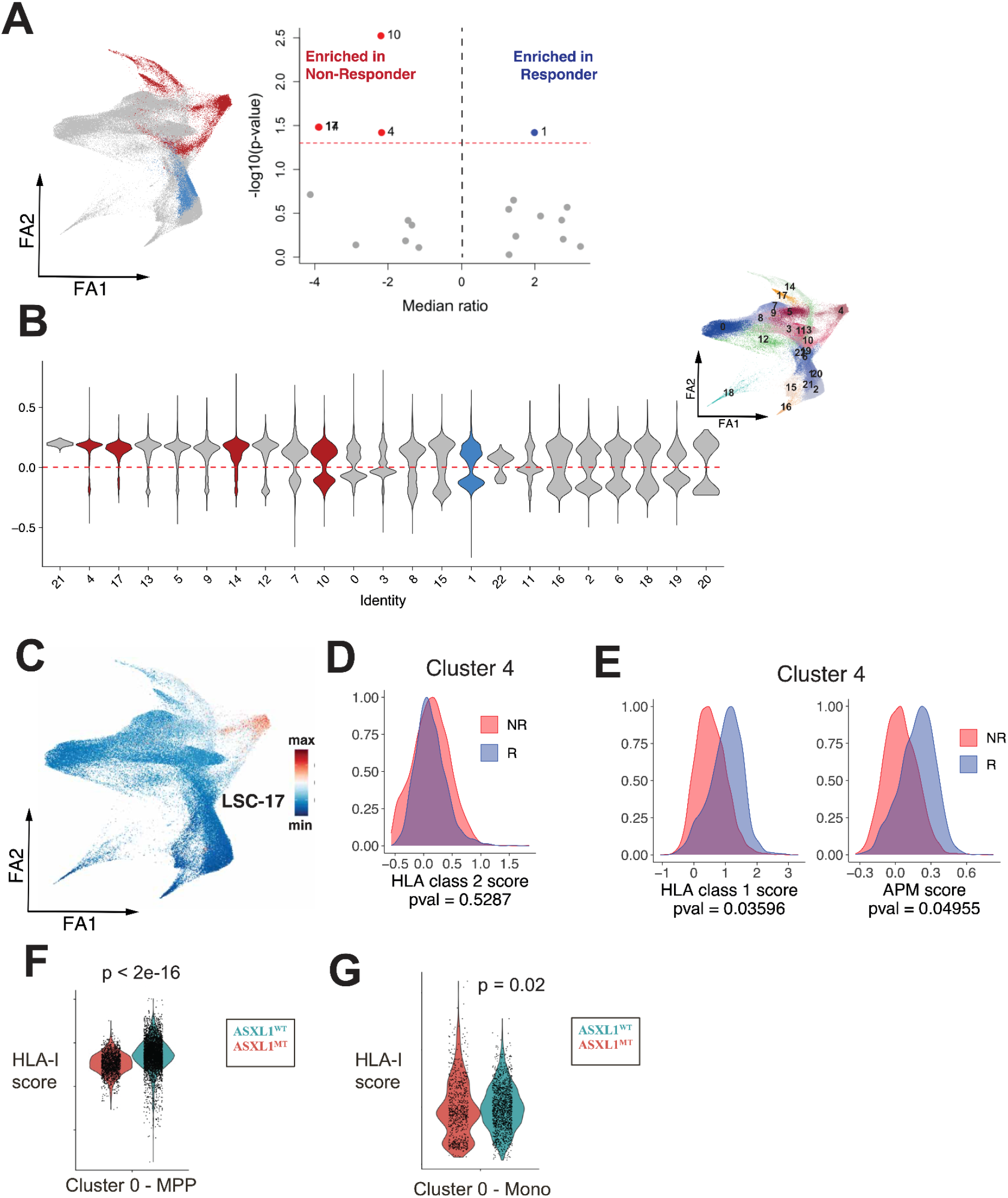
DLI resistance associates with HLA-1 suppression. A)Clusters differentially enriched across patients in non-responders vs responders. B) Clusters enriched in leukemic cells as assessed by donor/recipient score. C)SC17 score nominates cluster 4 as a cluster composed of leukemic stem cells. D)HLA-II is not differentially enriched in responders or nonresponders within cluster 4. E) HLA-I and antigen-processing machinery (APM) score within cluster 4 identifies differential expression of these genes across responders and nonresponders.Suppressed HLA-I within leukemic stem cell clusters from *ASXL1* mutationan external scRNA-seq AML cohort. G)Reduced accessibility of HLA-I genes within leukemic stem cell clusters in an external scATAC-seq CMML cohort.

We next sought to determine if these clusters were enriched in recipient cells utilising our D/R-score. This demonstrated that multiple clusters were enriched in recipient cells (Figure 3b). The four clusters enriched in non-responders were composed of varying proportions of recipient cells, with the composition of the GMP cluster being distributed across donor and recipient cells. In contrast, the HSC-like cluster was enriched in recipient cells, consistent with the known existence of BCR-ABL mutant CD34+ progenitors in CML.^27,28^

Given that CD34+ BCR-ABL mutant CD34+ progenitors are known to be a source of CML leukemic stem cells (LSC) ^11,27–31^ we sought to further characterize this HSC-like cluster. Using multiple transcriptional signatures for LSC (Figure 3c), we identified that this cluster demonstrated transcriptional evidence of being bona-fide LSC. Next, we sought to investigate potential immune escape mechanisms within this cluster. Given the known role for HLA dysregulation at leukemic relapse following alloSCT^32–36^ we examined HLA-I and HLA-II gene expression. Surprisingly, we did not identify alterations in HLA-II expression as has been previously reported during AML relapse after alloSCT^32,35,36^ (Figure 3d). In contrast, we identified decreased HLA-I (HLA-A,B,C) expression in DLI-NR (Figure 3e). These findings suggested that *ASXL1*-mutations may mediate immune escape through suppression of HLA-I expression.

### Validation in additional *ASXL1* mutant cohorts

To test this hypothesis in additional myeloid malignancies, we probed existing datasets in *ASXL1* mutant myeloid malignancies. A data set of nine patients with acute myeloid leukemia (AML)^10^ demonstrated that in a cluster with enrichment in LSC-signatures, cells from patients with *ASXL1*-mutant leukemia demonstrated significantly decreased HLA-I (HLA-A,B,B2M) (Figure 3f). Separately in a cohort comprised of five patients with chronic monomyelocytic leukemia (CMML)^9^, we utilised single-cell chromatin accessibility sequencing (scATAC-seq) data to identify clusters with higher accessibility of LSC-related genes and evaluated the accessibility of HLA-I genes. Consistent with the lower dynamic range of scATAC-seq^37^, there was decreased accessibility of HLA-I (HLA-A,B,C,B2M) in *ASXL1*-mutant cases, though this difference was less than in the scAML cohort (Figure 3g).

### Functional interrogation in leukemic cell lines

To functionally interrogate the contribution of *ASXL1* mutations to HLA-I suppression, we characterized K562, a pluripotent hematopoietic stem cell like cell line derived from a CML patient in blast crisis carrying an *ASXL1* mutation (Y591X)^38–43^ known to be HLA-I negative. We previously reported the creation of K562 isogenic cell lines in which the endogenous *ASXL1* Y591X mutation was corrected utilizing CRISPR-Cas9 (K562^m^)^44^ (Figure 4a). low cytometry demonstrated significantly increased cell surface protein expression of HLA-I, but not HLA-II, (Figure 4b, 4c) following *ASXL1* Y591 correction (K562^c^). Treatment with interferon-gamma (IFNγ) is known to potently increase cell surface HLA-I in K562,^45^ so we treated both cell lines with IFNγ. The K562^c^ cell line demonstrated significantly increased HLA-I compared to K562^m^ (Figure 4a,4b), though neither were able to increase HLA-II. To determine whether changes in HLA-I were due to increased transcription, we performed RNA-seq of the isogenic cell lines. K562^c^ demonstrated significantly increased HLA-I expression (Figure 4d).

**Figure 4.**
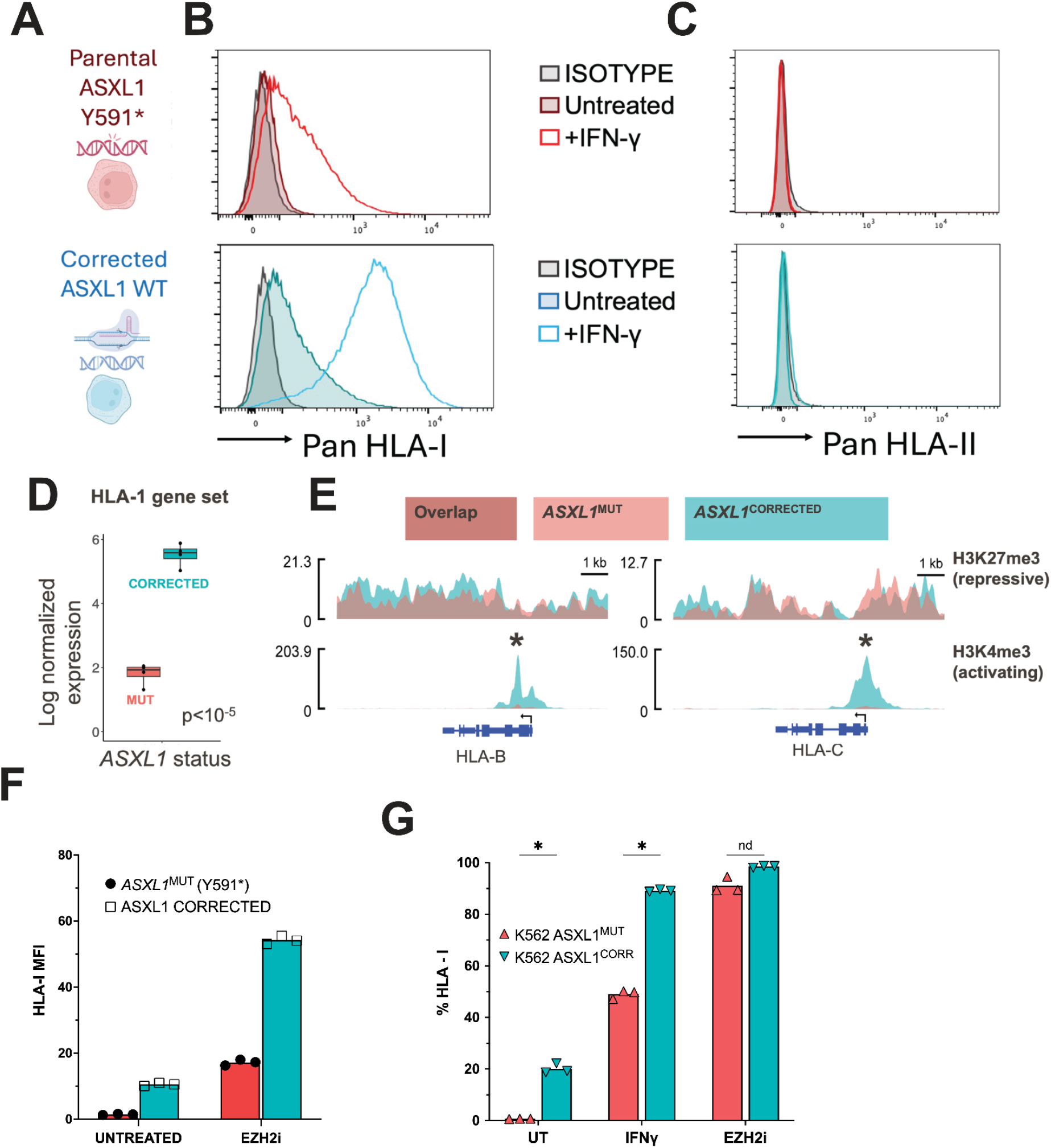
*ASXL1*^MUT^ represses HLA-1 surface protein expression through histone remodeling. A) Cartoon schematic of CRISPR corrected, isogenic cell lines. Flow cytometric measurements of B) HLA-I and C) HLA-II surface protein in the presence/absence of IFNg. D) RNA-expression of HLA-I genes in isogenic K562 cell lines. E) SparK plots of H3K4me3 and H3K27me3 at HLA-B and HLA-C genes following CUT&RUN profiling. F) Flow cytometry identification of HLA-I following 5 day treatment with EZH2i. G) Flow cytometry identification of HLA-I following 9 day treatment with EZH2i and 24-hour treatment with IFNγ treatment.

Since *ASXL1* is a known chromatin remodeler, we hypothesized that the changes in transcription may be driven by epigenetic alterations. We profiled H3K4me3 and H3K27me3 with CUT&RUN, as the presence of these marks at the same histone results in a state of bivalency that results in tightly regulated gene transcription. Moreover, *ASXL1* mutations have been implicated in altered H3K4me3 distribution across the genome. At both HLA-B and HLA-C, we did not detect alterations in H3K27me3, but did identify differentially enriched peaks at both HLA-B and HLA-C (Figure 4e).

H3K27me3 is deposited by the Polycomb Repressive Complex 2 (PRC2) through the enzymatically functional subunit, Enhancer of Zeste 2 (EZH2). Inhibitors of EZH2 (EZH2i) are being tested in multiple malignancies and are approved in follicular lymphoma. Given our finding of preserved H3K27me3 at HLA-genes in the isogenic cell lines, we hypothesized that treatment with EHZ2i would increase HLA in both cell lines. Consistent with this, we identified that treatment with EZH2i over 5 days can increase cell surface HLA in both K562^m^ and K562^c^ cell lines, with K562^c^ cell line achieving nearly university HLA-I positivity (Figure 4f). Following an extended treatment with EZH2i over nine days, both cell lines achieved nearly universal HLA-I expression (Figure 4g).

Given that truncating mutations in ASXL1 are known to enhance the activity of the Polycomb Repressive-Deubiquitinase complex (PR-DUB) in removing H2AK119Ub across the genome,^11,14,15,44^ we also performed H2AK119Ub CUT&RUN. We did not detect differences at HLA-genes, but we were able to confirm global reduction of H2AK119Ub (Figure 4h) consistent with previous reports.^44^

### *ASXL1* correction restores T-cell attack

We next sought to determine whether correcting *ASXL1*^m^ could restore T-cell targeting of K562 by T-cells. To do this, we developed a K562 T-cell recognition assay built upon our prior work generating antigen-specific T-cells^46^ (Figure 5a). Replacing peptide antigen with a 50%/50% mixture of irradiated IFNγ pre-treated K562^m^ and K562^c^ isogenic cell lines, we expanded T-cells with IL-7 (40ng/mL) and IL-2 (150IU) over the course of 14 days, before restimulating T-cells with a fresh 50%/50% mixture of irradiated IFNγ pre-treated K562^m^ and K562^c^ isogenic cell lines. After 21 days of total culture, T-cells were co-cultured with CellTrace™ Violet (CTV) labeled K562^m^ and K562^c^ over 18 hours before staining with viability dye and cell surface markers. Control T-cells were stimulated with anti-CD3 and anti-CD28 beads for 1 week prior to co-culture. K562 primed T-cells demonstrated preferential recognition of K562^c^ compared to control T-cells, while K562^m^ demonstrated no preferential recognition even at 10:1 E:T ratio (Figure 5b). Co-staining of CD3, CD8, and CD137 demonstrated upregulation of CD137 in K562 primed T-cells upon K562^c^ co-culture and not K562^m^ (Figure 5c).

**Figure 5.**
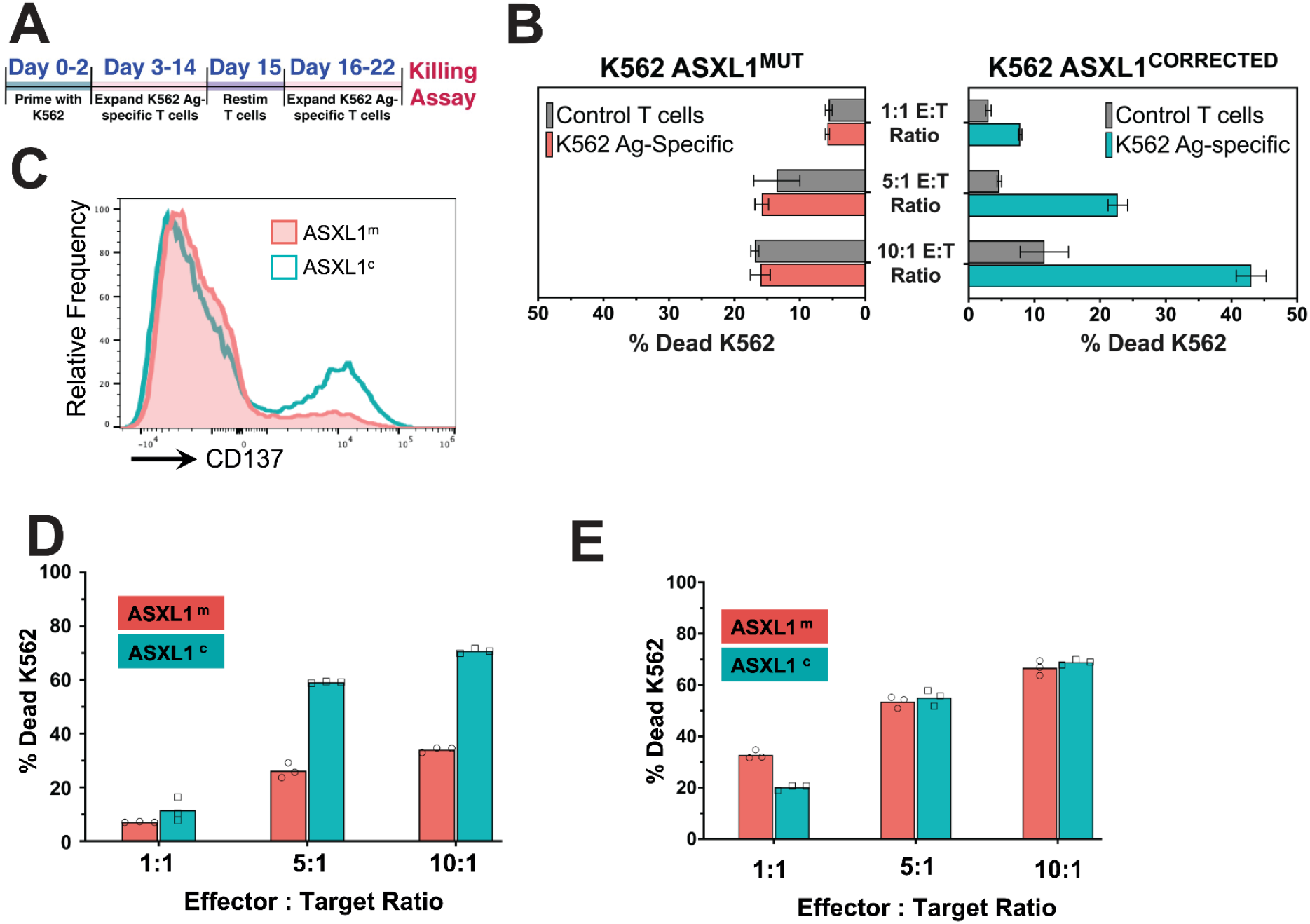
Correction of ASXL1 mutations or EZH2i pre-treatment restores T-cell attack. A) Experimental layout of anti-leukemic cell line T cell generation. B) Comparison of K562 killing by control or K562 primed T cell. C) Histogram demonstrating CD137 expression upon re-encounter with K562, either K562^MUT^ or K562^CORRECTED^. D) Persistent differences in K562 killing following IFNγ pretreatment. E) EZH2i pretreatment restores killing of K562^MUT^.

We then pretreated K562^m^ and K562^c^ with either IFNγ (24 hours) or EZH2i (5 days) prior to CTV labeling to determine whether these agents could enhance T-cell killing of K562^c^ and K562^m^. Pretreatment with IFNγ enhanced T-cell recognition of K562^c^ and enabled preferential T cell killing of K562^m^ compared to control T-cells, though primed T-cells killed K562^c^ much more readily than K562^m^ (Figure 5d). In contrast, pre-treatment with EZH2i enabled T-cell killing of K562^c^ and K562^m^ at similar levels, suggesting that EZH2i may be able to circumvent ASXL1^m^ suppression of T-cell recognition and killing (Figure 5e).

## Discussion

Post-SCT relapse is the greatest obstacle to SCT success, representing a failure of effective GvL. Understanding the pathways driving GvL evasion offer new opportunities to reinvigorate anti-leukemic immunity. Indeed, such insights may also impact anti-leukemic strategies in the SCT-naive setting as well. Using a well-annotated cohort of DLI treatment in the absence of concomitant immunomodulatory agents or chemotherapy and in the relatively simple genetic context of CML, we identified truncating mutations in the *ASXL1* oncogene as the genetic basis for DLI resistance. Integrative analysis with scRNA-seq data revealed HLA-I suppression as a key mechanistic insight into this oncogenetic circuit. Indeed, its relevance beyond the setting of alloSCT was highlighted by our analyses of publicly available scRNA- and ATAC-seq data of SCT-naive AML and CMML patients with similarly pathogenic *ASXL1* mutations. Mechanistically, we demonstrated that *ASXL1*^MUT^ repress HLA-I expression by decreasing the deposition of activating H3K4Me3 histone marks. This epigenetic activity decreases cell surface expression of the HLA-I protein, leading to CD8+ T cell quiescence and diminished cytotoxicity. Importantly, inhibition of the EZH2 pathway could overcome *ASXL1*^MUT^-mediated HLA-1 suppression and reinstate CD8+ K562-specific killing. Taken altogether, our findings identify a new oncogenetic pathway that governs the expression of HLA-I in myeloid malignancies. Additionally, we define how *ASXL1* mutations shape epigenetic marks that regulate transcription of HLA-genes. Moreover, we identify EZH2i as a potential agent, in contrast to IFNγ that can overcome *ASXL1* mediated HLA-I suppression. Finally, correcting *ASXL1* mutations or treatment with EZH2i can enable T cell killing of *ASXL1-*mutant leukemia.

## Methods

### Human subjects

Bone marrow (BM) biopsies were obtained pre- and post-DLI after relapse following alloSCT (or during remission following alloSCT) from patients enrolled in Dana-Farber Cancer Institute (DFCI) clinical trials (94-009, 95-011, 96-372, 96-022, and 96-277) between 1994-2001 that were approved by the DFCI Human Subjects Protection Committee. These studies were conducted in accordance with the Declaration of Helsinki; informed consent was obtained from the patients. Bone marrow mononuclear cells (BMMCs) were isolated via Ficoll-Hypaque density gradient centrifugation, cryopreserved with 10% dimethyl sulfoxide, and stored in vapor-phase liquid nitrogen until the time of sample processing.

### Cohort sample characteristics

All 17 patients had CML that was treated with CD8 T cell depleted alloSCT. Of these, 15 patients had CML relapse after alloSCT that was treated with CD8-depleted DLI, and 2 patients never had CML relapse and served as non-relapse controls. DLIs were infused at weekly intervals until the target cell dose was reached.

### Sample processing

Cryopreserved primary bone marrow mononuclear cells (BMMCs) were thawed on the day of sequencing at 37C and dispensed drop-wise into a warmed solution of 10% FBS, 10% DNaseI (StemCell Technologies, cat. No. 07900) in PBS. The cell suspension was centrifuged at 200 g for 10 minutes at room temperature. Viable cells were negatively selected using MACS Dead Cell Removal Kit (Miltenyi Biotec, cat. No. 130-090-101), running on MS columns to prevent sample loss. Collected live cells were resuspended in 0.04% BSA in PBS and diluted to a concentration of 1000 cells/uL. These cells were then divided into portions taken immediately for scRNA-seq. Bone marrow mesenchymal stromal cells (BM-MSCs) were expanded from thawed bone marrow vials in 10% FBS in DMEM, cultured as adherent cells, and isolated as live, CD45 negative expressing populations.

### WES data generation and preprocessing

For WES, standard protocols were used as previously described.^17^ Genomic DNA from all samples was isolated using DNeasy Blood & Tissue Kit (Qiagen). Tumor DNA was derived from thawed BMMCs or PBMCs as indicated; matched germline DNA derived from both donor (e.g. T cells expanded from post-SCT PBMCs or DLI products) and recipient (i.e. BM-MSCs) sources. DNA concentrations were measured using PicoGreen dsDNA Quantitation Reagent (Invitrogen). A minimum DNA concentration of 5 ng/l was required for sequencing. All Illumina sequencing libraries were created with the native DNA. The identities of all tumor and normal DNA samples were confirmed by mass spectrometric fingerprint genotyping of 95 common single nucleotide polymorphisms (SNPs) by Fluidigm Genotyping (Fluidigm). Sequencing libraries were constructed with the Illumina Rapid Capture Enrichment Kit. Pooled libraries were normalized to 2 nM and denatured using 0.2 N NaOH before sequencing. Flowcell cluster amplification and sequencing were performed according to the manufacturer’s protocols using either the HiSeq 2000 v3 or HiSeq 2500. Each run was a 76–base pair paired-end with a dual eight-base index barcode read. Standard quality control metrics, including error rates, percentage-passing filter reads, and total Gb produced, were used to characterize process performance before downstream analysis. Data analysis included de-multiplexing and data aggregation.

### Mutation calling

We used the Genome Analysis Toolkit (GATK) pipeline developed at the Broad Institute as previously described.^17^ Paired-end Illumina reads were aligned to the hg19 human genome reference using the Picard pipeline (https://software.broadinstitute.org/gatk/documentation/tooldocs/4.0.1.0/picard_fingerprint_CrosscheckFingerprints.php; https://software.broadinstitute.org/gatk/documentation/tooldocs/4.0.0.0/picard_analysis_CollectMultipleMetrics.php). Cross-sample contamination was assessed with the ContEst tool (5% threshold). Point mutations and indels were identified using MuTect and Strelka, followed by annotation using Oncotator. Possible artifacts due to orientation bias, germline variants, sequencing, and poor mapping were filtered using a variety of tools including Orientation Bias Filter, MAFPoNFilter, and RealignmentFilter. For pre-SCT samples, somatic mutations were called using recipient fibroblast as paired normal control samples. For post-SCT samples, we first call somatic mutations separately using either recipient fibroblast or donor PBMCs/DLI products as paried normal control samples. Then we took the intersection of the two sets as the somatic mutations called somatic mutations. We further filtered out all somatic variants with allele frequency > 0.001 in genomeAD (https://gnomad.broadinstitute.org/) using Variant Effect Predictor (VEP v98). All somatic alterations thus identified were further manually inspected in IGV, and apparent false positives were removed.

### Flow Cytometry

1,000,000 isogenic K562 cells were cultured in 10% IMDM media. Cells were treated with EZH2i (EPZ011989, Selleckchem, 3uM) or IFNg (Biolegend, 50ng/mL) for the duration indicated in the figure legend. Cells were washed in PBS prior to staining with Live-Dead NIR followed by surface staining for HLA-I (W6/32) either in PE or R-718 (all BD Biosciences) and HLA-II (Tu39) in BV711, BV421, or PE (all BD Biosciences).

### RNA-seq

RNA was isolated from 1,000,000 isogenic K562 cells using RNeasy Kits and library preparation, and RNA-seq was performed at the MD Anderson Advanced Technology Genomics Core (ATGC) using standard protocols. For IFNg treated groups, K562 was cultured with IFNg (50ng/mL, Biolegend) for 24 hours prior to RNA isolation. A complementary DNA (cDNA) library was prepared from poly-A–selected RNA and sequenced on an Illumina platform. Paired-end transcriptome reads were mapped to the GRCh38 reference genome using STAR. Differential expression analyses were conducted using DESeq2 R package v.1.38.3. Differentially expressed genes were identified using a cutoff of FDR P ≤ 0.25. GSEA analysis was performed using fgsea (v 1.24.0).

### CUT&RUN

CUT&RUN chromatin sequencing for H3K4me3, H3K27me3, and H2AK119Ub isogenic K562 cell lines was performed according to published protocols.^47^ 200,000 cells were washed twice in CUT&RUN wash buffer (20 mmol/L HEPES, 150 mmol/L NaCl, 0.5 mmol/L Spermidine, supplemented with Roche EDTA-free cOmplete Protease Inhibitor). Next, activated Concanavalin A–coated magnetic beads (BioMagPlus Concanavalin A, Bang Laboratories) were added to the cell suspension, and cells were allowed to bind to the beads for 10 minutes at room temperature under gentle agitation. After washing, the cells/bead mixture was resuspended in antibody buffer (consisting of wash buffer supplemented with 0.1% digitonin, 2 mmol/L EDTA, and 0.5% BSA) and appropriate histone antibodies or IgG control (Clone DA1E, Cell Signaling Technologies) at a 1:50 dilution and incubated with gentle agitation at 4°C overnight. After washing twice to remove unbound antibody, protein A/G-MNase fusion protein (Epicypher) was added to the cell/bead mixture and allowed to bind for 2 hour at 4°C. Cells were washed twice to remove unbound antibodies, resuspended in wash buffer supplemented with 0.1% digitonin, and allowed to cool to 0°C in a chilled ice block before adding CaCl_2_ to a final concentration of 2 mmol/L to initiate cleavage. After 30 minutes, the reaction was quenched with EDTA and EGTA, and cleaved fragments were allowed to be released into the supernatant by incubating the cells at 37°C for 30 minutes. Fragments were analysed on a Bioanlyzer before clean up and library preparation performed by the MD Anderson Epigenetics Profiling Core. We used nf-core/cutandrun (https://nf-co.re/cutandrun/3.2.2/) bioinformatic analysis pipeline to process raw Cut and Run data. Briefly, after trimming of adapters using TrimGalore (**https://doi.org/10.5281/zenodo.5127899**), reads were mapped to GRCh38 using Bowtie2 (PMID: 22388286). Alignments in bam format were then converted to bigwig files using bedtools (PMID: 20110278) and bedGraphToBigWig for visualization in IGV. H3K4me3, H3K4me2, H3K36me3, H2AUb119, and H3K27me3 occupancy peaks were called using SEACR (PMID: 31300027) and consensus peaks were merged using bedtools. Peak-based quality control including fingerprint, PCA, and Correlation heatmap plots was carried out using deepTools (PMID:27079975). Differential abundance analysis comparing ASXL1 WT and MT for each histone mark was done using DESeq2 (PMID: 25516281) and peaks were annotated using ChIPpeakAnno (PMID: 20459804).

### T-cell killing assays

T-cells were generated according to the previously published protocol^46^ with the following modifications. 1. IFNg pre-treated (50ng/mL, Biolegend, 24-hours) irradiated K562 cell lines were used for an antigen source and added at 1E6 per 2E6 PBMC on Day 1. 2. LPS was not used. Instead, IFNg (50ng/mL, Biolegend), was added. 3. 40ng/mL IL7 (Miltenyi Biotec) and 150IU/mL IL2 (Peprotech) were used to expand T-cells starting on Day 2. For killing assays, non-irradiated K562 cells were labelled with CellTrace Violet according to manufacturers instructions and plated with antigen-specific T-cells at the indicated E:T ratios in 10% FBS IMDM and cultured for 18 hours. For the pre-treated groups (IFNg & EZH2i), isogenic K562 cell lines were pre-treated with IFNg (50ng/mL, Biolegend, 24 hours) or EZH2i (EPZ011989, Selleckchem, 3uM, 5 days). Following co-culture, cells were stained with Live-Dead NIR followed by CD3 PE-Fire 640 (Biolegend), CD8 BV650 (Biolegend), CD4 BB515 (BD Biosciences), CD56 BUV737 (BD Biosciences), HLA-I R-718 (BD Biosciences), and CD137 APC (BD Biosciences) and read on a Cytek Aurora. Dead K562 was calculated as (#L-D NIR+/#CTV+)*100 and performed in triplicate. Comparisons were performed with two-tailed T-test.

## Acknowledgements

SL is supported by NCI R50 Research Specialist Award (R50CA251956). S. A. Shukla is supported by the Cancer Prevention and Research Institute of Texas (CPRIT) award RR220009 and is a CPRIT Scholar in Cancer Research. C.J.W. is supported in part by National Institutes of Health, National Cancer Institute grants P01CA229092, the LLS SCOR-22937-22 and is a member of the Parker Institute for Cancer Immunotherapy at DFCI. Her work is also supported, in part, by the Parker Institute for Cancer Immunotherapy. C.J.W. is also the Lavine Family Chair for Preventative Cancer Therapies at DFCI. P.B. is a CPRIT Scholar in Cancer Research and an Andrew Sabin Family Foundation Fellow at The University of Texas MD Anderson Cancer Center and is supported by an Amy Strelzer Manasevit Scholar Award from the Be The Match Foundation, by the NCI grant 1K08CA248458-01, and Break Through Cancer funding.

